# An integrated map of genetic variation from 1,062 wheat genomes

**DOI:** 10.1101/2023.03.31.535022

**Authors:** Aoyue Bi, Daxing Xu, Lipeng Kang, Yafei Guo, Xinyue Song, Xuebo Zhao, Jijin Zhang, Zhiliang Zhang, Yiwen Li, Changbin Yin, Jing Wang, Fei Lu

**Affiliations:** State Key Laboratory of Plant Cell and Chromosome Engineering, Institute of Genetics and Developmental Biology, The Innovative Academy of Seed Design, Chinese Academy of Sciences, Beijing, China; University of Chinese Academy of Sciences, Beijing 100039, China; CAS-JIC Centre of Excellence for Plant and Microbial Science (CEPAMS), Institute of Genetics and Developmental Biology, Chinese Academy of Sciences, Beijing, China

**Author notes:** Corresponding author (F.L.). These authors contributed equally to this work (Aoyue Bi, Daxing Xu).

## Abstract

The construction of a high-quality wheat genome variation map is important to wheat genetic studies and breeding. In this study, we developed the second-generation whole-genome genetic variation map of wheat (VMap 2.0) by integrating whole-genome sequencing data of 1,062 diverse wheat accessions from 20 species/subspecies. VMap 2.0 contains 195.96 million single nucleotide polymorphisms (SNPs), 2.22 million insertions, and 4.72 million deletions, achieving a high density of variation map in which one variant exists in every 73 base pairs on average. By systematically analyzing the phylogenetic relationships and genetic diversity of tetraploid wheat, hexaploid wheat, and diploid goatgrass (*Aegilops tauschii*), we found that the genetic diversity of wild emmer wheat was 2.4 times higher than that of common wheat. In contrast, the genetic diversity of diploid goatgrass is 7.8 times higher than the D subgenome of hexaploid wheat. With the high-density genetic variations, VMap 2.0 is anticipated to facilitate high-resolution trait dissection and expedite prediction-based breeding of wheat.

## Main text

High-density genetic variation map is a prerequisite for high-resolution trait mapping and precision breeding. With the development of genomic and computational technology, it has become a foundation for crop improvement. Exciting opportunities are emerging for the integration of whole-genome genetic variations from numerous biological materials worldwide. This will advance our understanding of crop biological processes, providing greater impetus for translating laboratory results to the field ^1^.

Bread wheat (*Triticum aestivum*, 2*n* = 6*x* = 42, AABBDD) is an essential staple crop, accounting for ∼20% of calories and protein consumed by humans globally ^2^. To develop improved wheat varieties, it is necessary to understand the genetic variation underlying important traits such as yield, drought tolerance, and disease resistance. To date, genetic studies of wheat populations have primarily been focused on the phylogenetic relationship, diversity and introgression ^3–8^, such studies have yielded valuable insight into wheat evolution. However, adaptive evolution from the perspective of functional variation has not yet been fully understood. The previously released wheat VMap 1.0 and VMap 1.1 were constructed with the whole-genome sequencing data of 414 and 795 accessions of bread wheat and its wild relatives, respectively ^4,8^. Since then, more wheat samples were sequenced with high depth, a more comprehensive variation map would help discover more functional variations and accelerate genomic and genetic research in wheat.

Here, we developed a high-density genetic variation map by collecting whole-genome sequencing data of 1,062 diverse wheat accessions (306 accessions are newly sequenced and 756 accessions are from previous studies ^3,4,8–11^, including hexaploid and tetraploid wheat, as well as diploid related species sampled around the globe. We aim to develop tools and resources to facilitate the use of this genetic variation map by wheat breeders, and characterize functional variations across ploidy levels, geographic distributions, and breeding status.

### High-quality variation set of the VMap 2.0

We aggregated diverse individual genomes with whole-genome resequencing data to characterize the functional variation in wheat populations (Supplementary Tables 1 and 2). The complete set of 1,062 genomes from 89 countries includes 212 tetraploid wheat, 814 hexaploid wheat, and 36 diploid *Ae. tauschii* (AT), of which subpopulations including wild emmer (WE, *n* = 91), domesticated emmer (DE, *n* = 52), free-threshing tetraploids (FTT, *n* = 50), other tetraploids (*n* = 19), landrace (LR, *n* = 306), cultivar (CL, *n* = 223) and other hexaploids (*n* = 285) (Supplementary Table 3). All these taxa were used to construct the whole-genome genetic variation map 2.0 of wheat (VMap 2.0), of which 306 taxa were newly sequenced, generating 469,864,946,476 pair-end reads and had average coverage of 10× (Supplementary Table 4). Each sequencing data set was uniformly aligned to the reference genome IWGSC RefSeq v1.0 ^12^ and processed by FastCall 2 for SNP discovery and genotyping (Fig. 1). However, considering the false positive variants probably caused by read mapping, an ‘accessible reliable genome’ was required to remove these artifacts ^13^. We developed the PopDepth pipeline for filtering regions of the reference genome with ambiguously placed reads and unreliable alignments. We then obtained a high-quality position database with reliable alignment to the correct genome location.

**Fig. 1.**
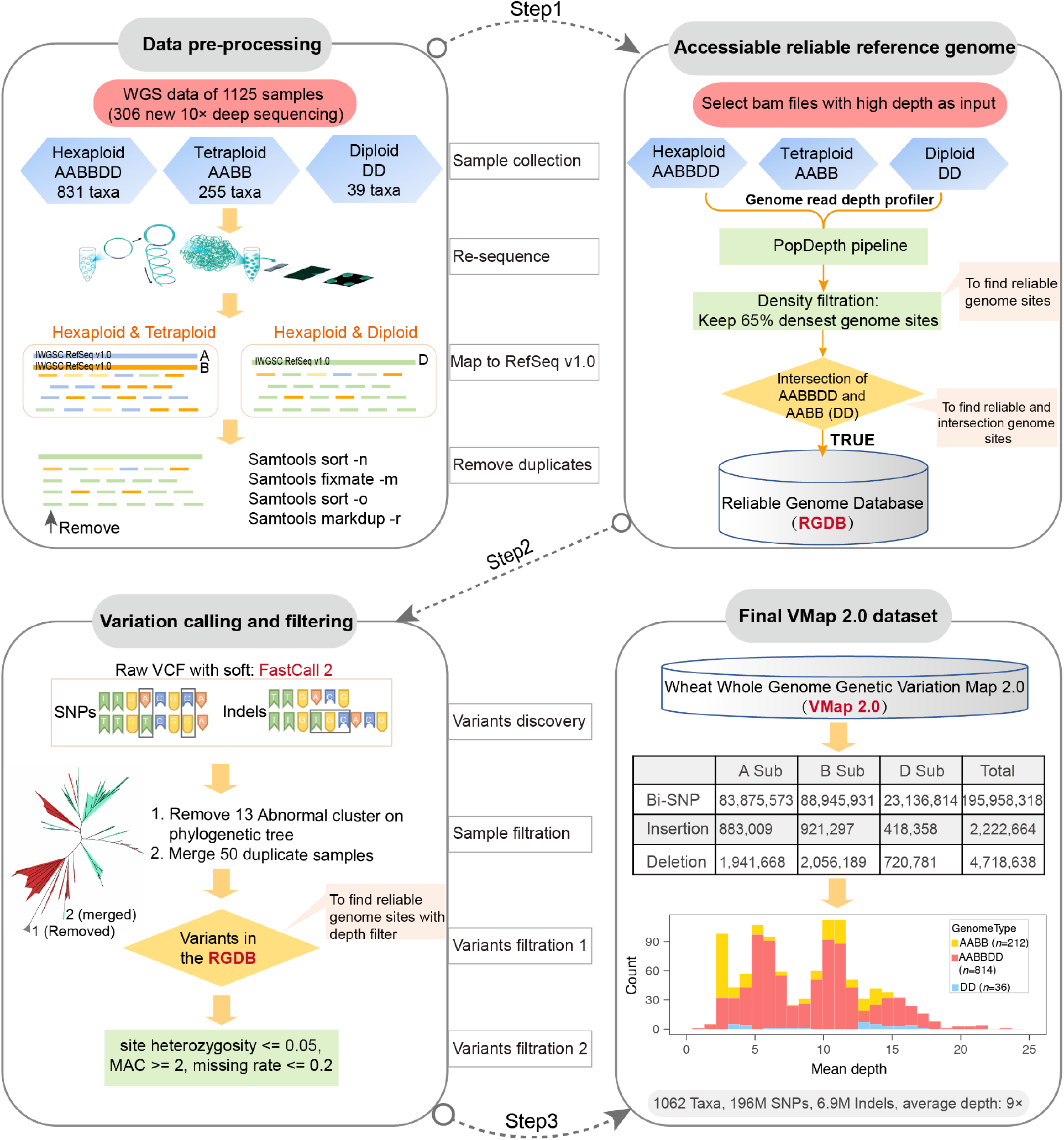
Overview of VMap 2.0 data processing workflow. Flowchart describing the process of samples and variants. First, 1,125 wheat and *Ae*.*tauschii* samples were collected and performed data pre-processing: BAM files with no duplicates are obtained through whole-genome resequencing and reference genome alignment. Second, as the accessible reliable reference genome make it better to call an accurate genetic variation set, we developed a new pipeline called “PopDepth” and then an initial depth filtration was performed to get the high-confidence genome variants using RGDB (reliable genome database). Subsequently, variants discovery was performed using FastCall pipeline. Sample duplication and data-quality were then checked using phylogenetic tree analysis. Finally, the wheat whole-genome genetic variation map 2.0 (VMap 2.0) were generated in further MAF and missing rate filtration. A total of 196 million SNPs, 2.2 millions insertion and 4.7 millions deletion were deposited in the wheat genomic database. The pipelines used in the study were released on https://github.com/PlantGeneticsLab/TIGER.

The final VMap 2.0 dataset was generated after the stringent sample quality control by site and taxon level, comprising 195,958,318 high-quality biallelic SNPs, 2,222,664 insertions and 4,718,638 deletions (Fig. 2a, Supplementary Fig. 1 and Supplementary Table 5). This variation data set contains an average of one variant every 50 and 100 bases relative to AB and D subgenomes (Fig. 2b). Of the total SNPs, 64%, 86%, and 60% in tetraploid, hexaploid and diploid were rare with minor alleles frequency less than 0.05, respectively (Fig. 2c). Nearly 48.8% and 21.8% of SNPs were population-specific among ‘tetraploid *vs*. hexaploid’ and ‘hexaploid *vs. Ae. tauschii*’ groups, respectively. The scores of identity-by-state (IBS) indicated the genetic distance to hexaploid Chinese Spring (Fig. 2d). Compared with wild emmer and domesticated emmer, free-threshing tetraploid is closer to Chinese spring (Fig. 3). After applying strict quality control at the sample level, the average depth of each variation site was 9.16× for hexaploid wheat, 7.45× for tetraploid wheat, and 11× for diploid *Ae. tauschii*. The variance of each variation site was also minimal, indicating that the filtered reads had a uniform comparison depth and met the expected filtering standards. Individual heterozygosity analysis revealed that hexaploid wheat had lower heterozygosity (mean value: 0.0065) than tetraploid wheat (mean value: 0.015), while *Ae. tauschii* had the highest heterozygosity (mean value: 0.017) (Fig. 4a). As the sequencing depth was high in this study (average: 9×), the genotype missing rate in most samples was below 0.05 (Fig. 4b). The sequencing depth of some tetraploid samples was 3-4×, resulting in a slightly higher missing rate (0.05-0.1). Some missing genotypes may also be caused by structural variation ^14^. Using the reference accession as a control, we detected the error rate of the variant calling of 6.73 × 10^−6^, which is lower than previous studies ^4,8,15^.

**Fig. 2.**
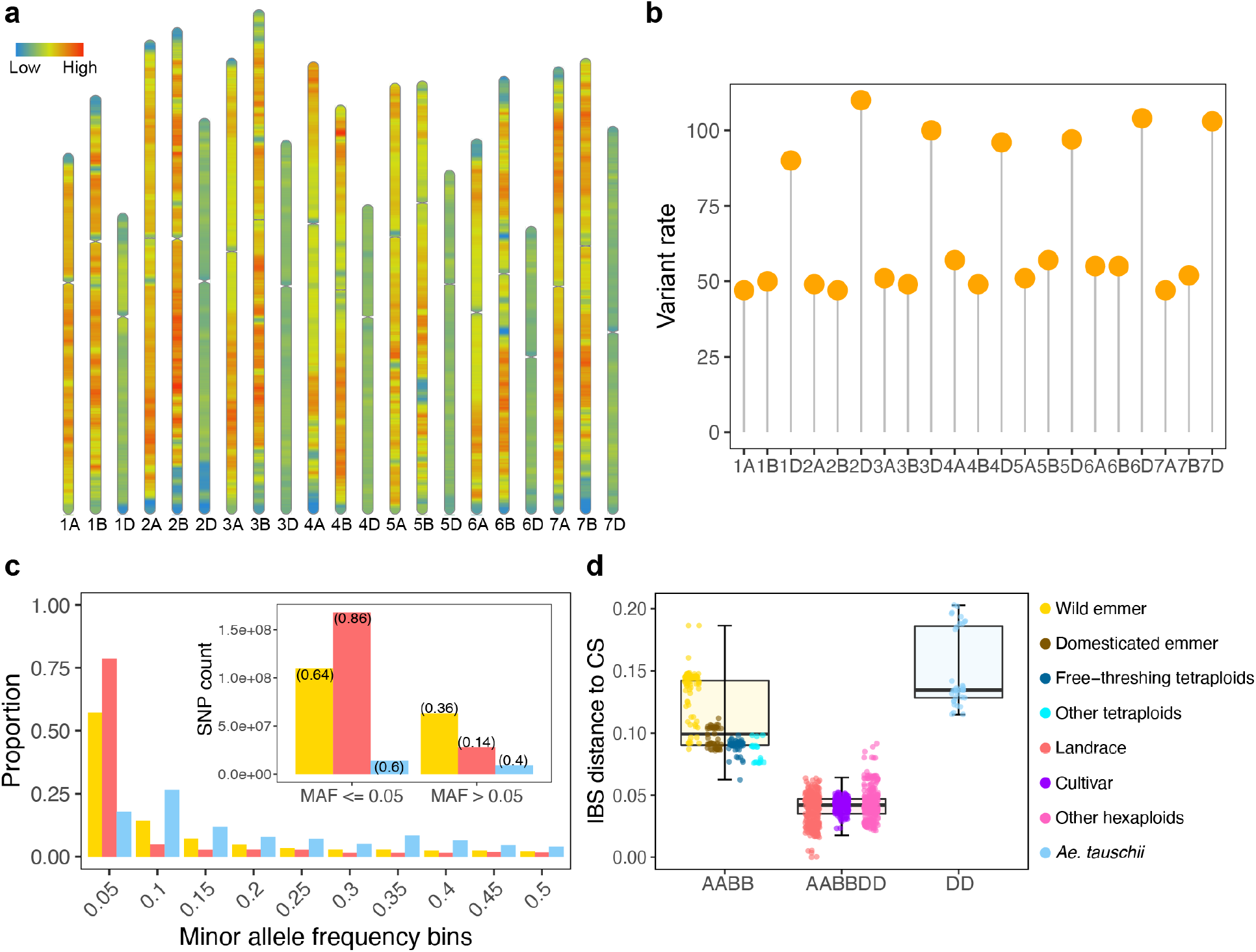
Variant discovery of VMap 2.0 at large sample size. **a**, Ideogram showing the high-density variation across the 21 chromosomes of VMap 2.0. Each chromosome was divided into bins of 10 Mb, with a sliding window of 1 Mb. **b**, Variant rate calculation across subgenomes. **c**, Minor allele frequency (MAF) distribution in hexaploid wheat (indian red), tetraploid wheat (yellow) and *Ae. tauschii* (sky blue) populations, the barplot inside showing SNP count in common and rare variants group. **d**, Boxplot showing the identity-by-state (IBS) distance to reference genome IWGSC RefSeq v1.0 in different ploidy populations. Each point on the boxplot represents a different individual colored by subpopulations.

**Fig. 3.**
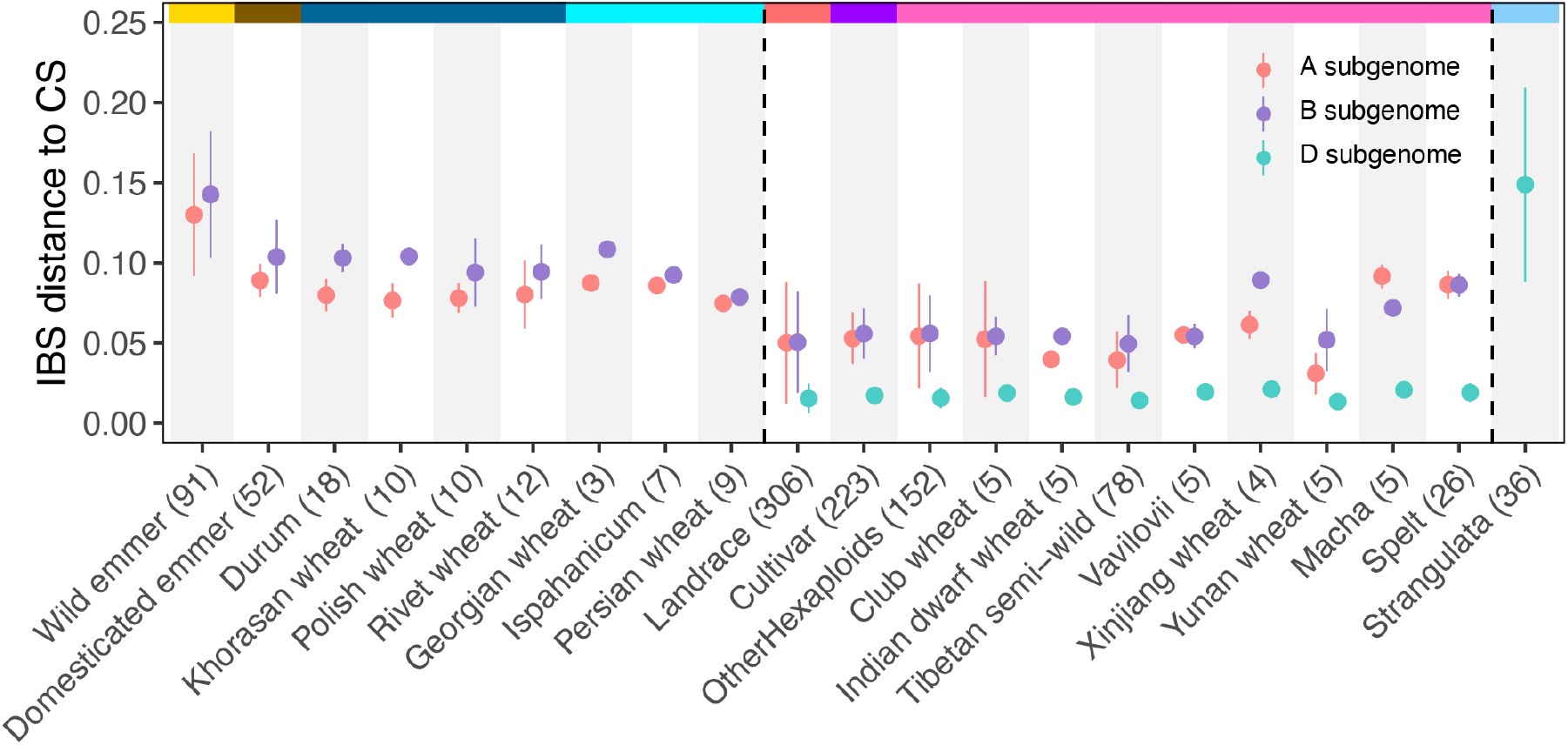
The mean IBS-distance in all subpopulations grouped by subgenomes (*n* = 1,062). Error bars represent the mean plus or minus the standard deviation.

**Fig. 4.**
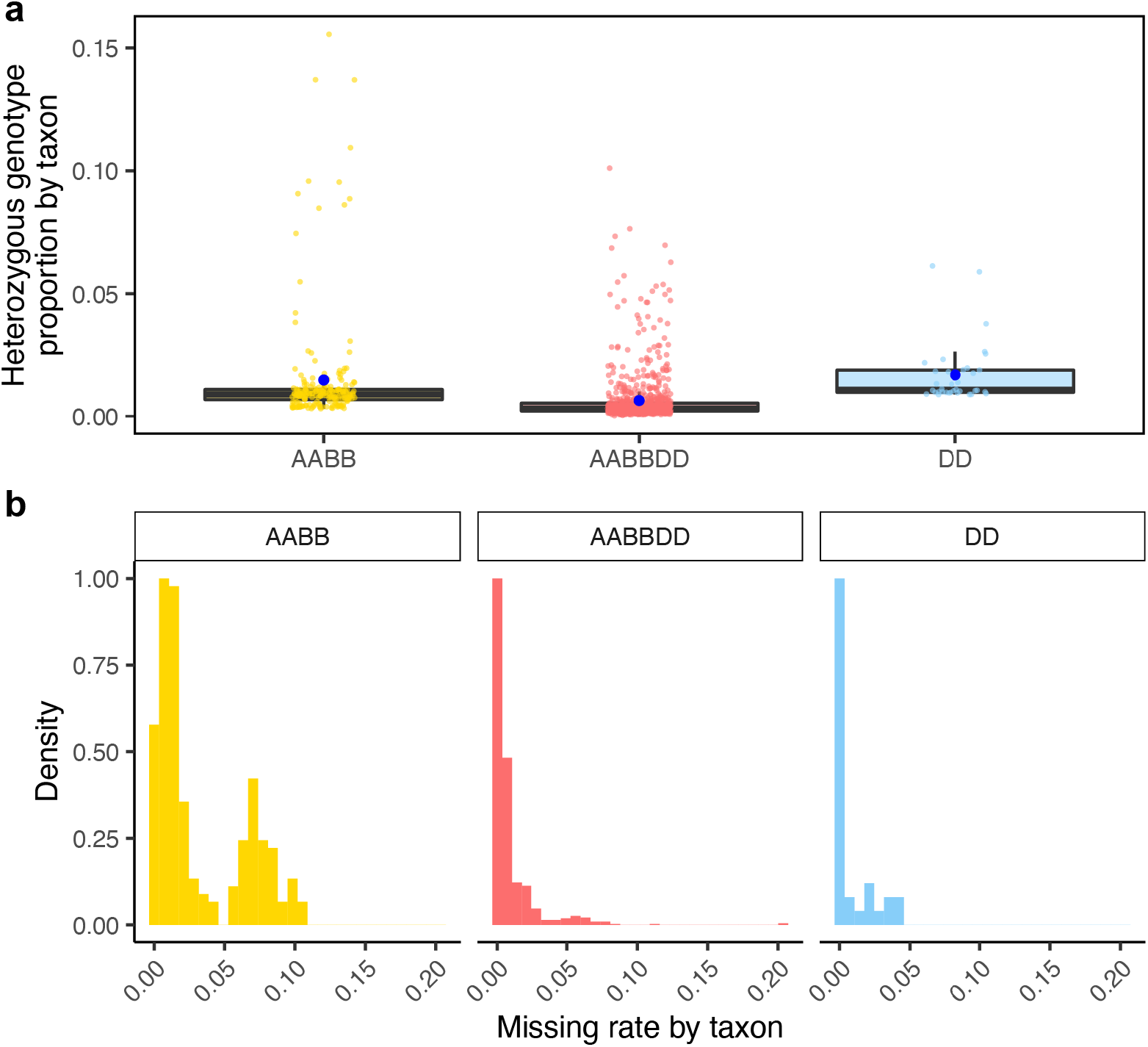
Sample quality control by taxon level of VMap 2.0. **a**, Boxplot showing the heterozygosity of individuals in different ploidy populations. Each point on the boxplot represents one taxon. The centerline of each box plot indicates the median and the lower and upper hinges indicate the 25th and 75th percentiles, respectively. The vertical line of each boxplot extends to 1.5× the interquartile range from each hinge. **b**, Histogram showing the missing rate of different ploidy species. The two double peaks in the tetraploid represent different batches of samples, in which the newly resequencing data has the lower missing rate because of the 10× coverage.

### The abundance of putative functional variants in gene region

The VMap 2.0 Gene Annotation Database (GADB) contained a total of 2,672,838 SNPs, which represented 1.36% of the whole-genome. Within GADB, the CDS region, intron region, 3’UTR, and 5’UTR accounted for 665,844, 1,693,079, 254,881, and 59,034 SNPs, respectively. These SNPs correspond to proportions of 24.91%, 63.34%, 9.54%, and 2.21% (Supplementary Fig. 2 and 3, Supplementary Table 6). Among the SNPs in the coding region, there were 286,341 synonymous mutations and 367,605 non-synonymous mutations, representing proportions of 10.71% and 13.75%, respectively. We investigated the relationship between sample size and variant detection rate, and observed that nonsense mutations were less frequent than synonymous and nonsynonymous mutations (Fig. 5 and Supplementary Fig. 4). The influence of purifying selection means that most functional mutations occurred at lower frequencies and were harder to detect, which was in line with findings from human genome sequencing research ^16,17^. Our study highlighted that the current sample size is insufficient to achieve full mutation saturation of the wheat genome, and suggested that there may be many rare functional variants still waiting to be uncovered within the wheat populations.

**Fig. 5.**
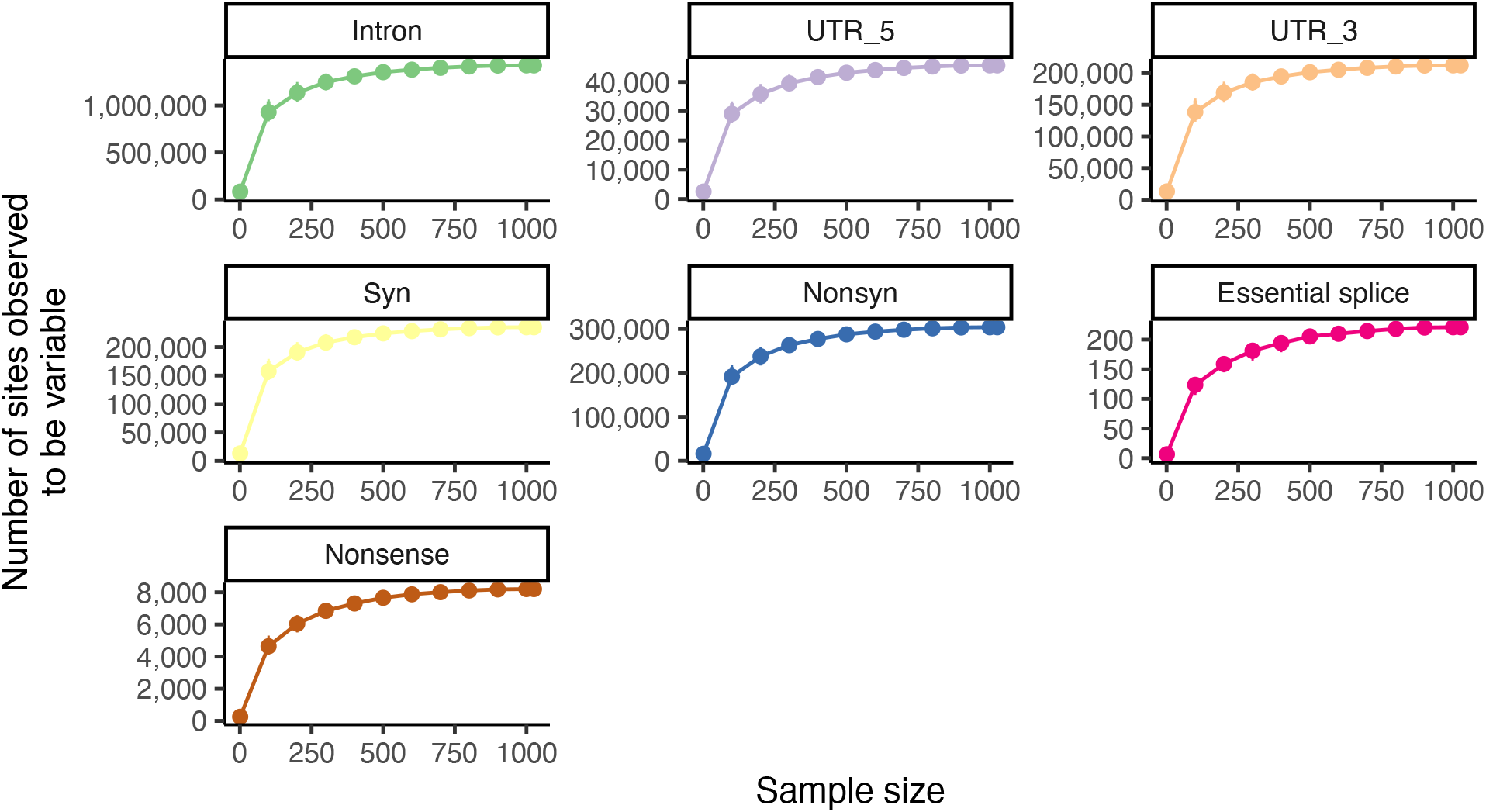
The total number of variants observed as a function of sample size in the AB subgenome.

### Population analysis of 1,062 wheat accessions

The principal component (PCA) analysis and phylogenetic inferences revealed a close relationship between *Triticum aestivum* and *Triticum turgidum* (Fig. 6 and Supplementary Fig. 5). A robust phylogenetic tree is an indispensable interpretive framework for our study of evolutionary relationships among wheat and its ancestors. Here we adopt the two-step phylogenetic method ^18^. First, we used barley (*Hordeum vulgare*) as an outgroup and constructed the neighbor-joining (NJ) tree to capture the most recent common ancestor (MRCA) of wild emmer-PI466987. Then we randomly sampled 230,000 bi-SNPs on the AB subgenomes of VMap 2.0 with a minor allele frequency (MAF) > 0.1 and reconstructed a comprehensive and informative phylogenetic tree rooted by the primitive wild emmer as ancestral state (Supplementary Fig. 5). In agreement with previous studies on the *Triticum* species tree ^4,7^, the whole-genome tree based on the AB lineage informed a chronological and reticulate evolutionary relationships, providing insights into the two-stage domestication process of tetraploid wheat. This process began with the domestication of Wild emmer wheat (*T. turgidum* subsp. *dicoccoides*, AABB) into cultivated emmer (*T. turgidum* subsp. *dicoccon*), which had a non-brittle rachis and was easier to harvest. Later, free-threshing tetraploid wheat, especially *T. durum*, evolved from hulled genotypes and is now widely grown. Finally, hexaploid bread wheat speciated through extensive genetic exchange (introgression and hybridization) between tetraploid emmer wheat (*T. turgidum*; AABB) and *Ae. tauschii* (DD) ^2,19–22^. Also, we observed that the clade representing hexaploid bread wheat is mainly composed of the Asian and European genetic groups. Among these, 93.27% (208/223) of cultivars were clustered on the European clade (Supplementary Fig. 5). Additionally, we found that the fixation index (*F*ST) between wild emmer and domesticated emmer in the A subgenome was greater (*F*ST = 0.265) than that between domesticated emmer and free-threshing tetraploid (*F*ST = 0.241) (Supplementary Fig. 6). Moreover, the genetic differentiation between hexaploid landraces and cultivars was much smaller than that between tetraploids, with *F*ST values of 0.061 and 0.041 in the A and B subgenomes, respectively (Supplementary Fig. 6). The average genetic diversity of wild emmer, domesticated emmer and free-threshing tetraploids wheat was 2.08 × 10^−3^, 1.38 × 10^−3^ and 0.94 × 10^−3^, respectively, which were 2.44, 1.63 and 1.1 times the hexaploid wheat (Supplementary Fig. 7). However, the mean genetic diversity of *Ae. tauschii* and hexaploid wheat’s D subgenome were 0.71 × 10^−3^ and 0.09 × 10^−3^, respectively, representing a 7.8-fold difference (Supplementary Fig. 7).

**Fig. 6.**
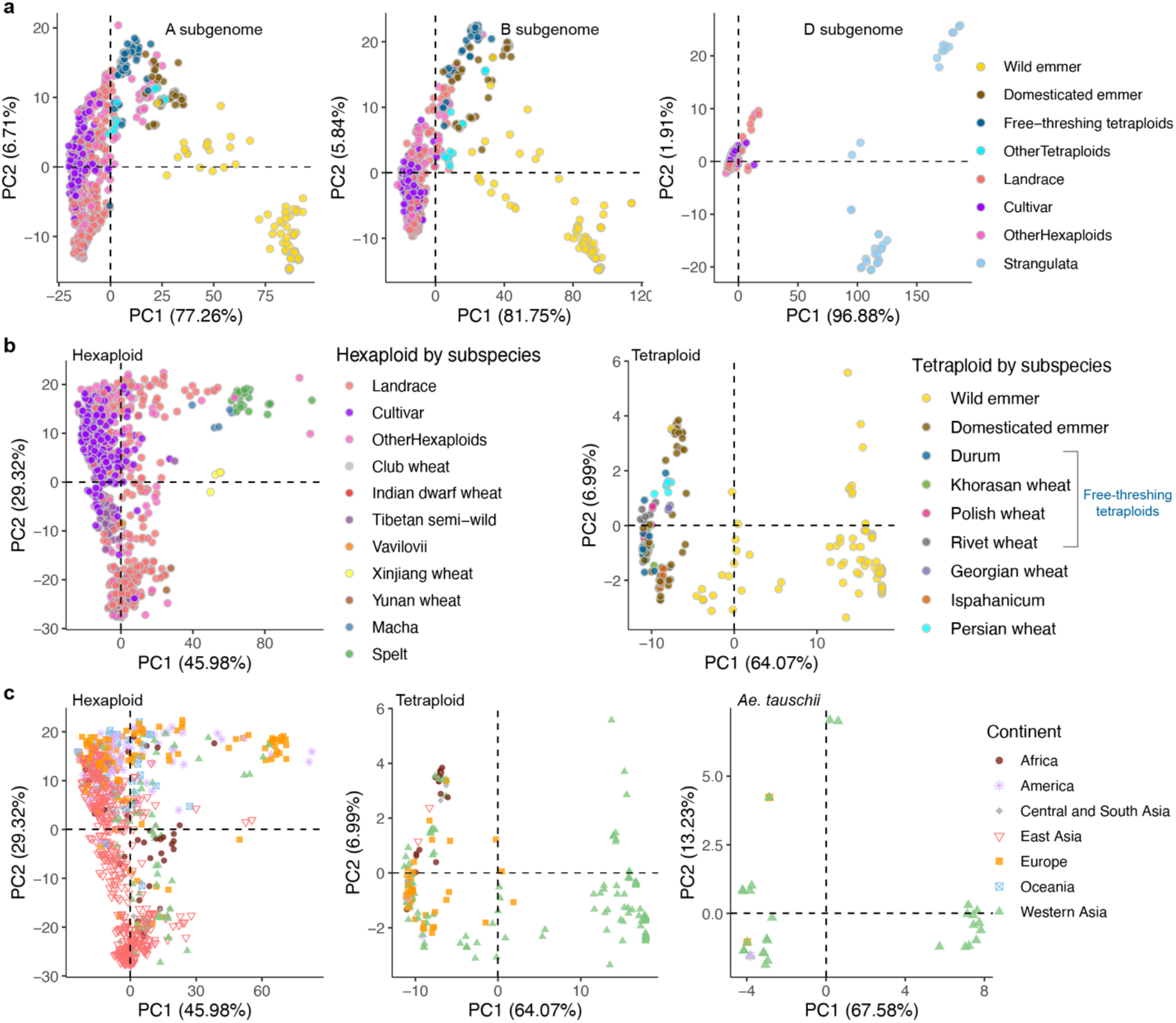
PCA of 1,062 diverse accessions on VMap 2.0. **a**, PCA of 1,026 taxa in the A and B subgenome as well as 850 taxa in the D subgenome. In the VMap 2.0, the AB subgenomes contains 814 hexaploid and 212 tetraploid wheat, while the D subgenome contains 814 hexaploid wheat and 36 *Ae. tauschii*. **b**, PCA of hexaploid and tetraploid wheat separately, the colored points correspond to the subspecies. **c**, PCA of hexaploid, tetraploid wheat and *Ae. tauschii*, the colored and shaped points correspond to the continents.

## Discussion

Crop improvement through efficient genome editing requires the identification of functional variants ^23^. Advancements in sequencing technologies and computational tools have enabled prediction of functional variation in wheat and its wild relatives. In this study, we constructed a whole-genome genetic variation map 2.0 with 1,062 wheat accessions and depicted a genome-wide landscape at a single base-pair resolution. By prioritizing functional variants based on their biological impact, we have built a platform for rapid mapping causative variation in wheat across populations, which may contribute to improvement of the trait prediction accuracy in wheat.

## Acknowledgments

This work was supported by the National Natural Science Foundation of China (32225038, 31970631) and Strategic Priority Research Program (A) of the Chinese Academy of Sciences (XDA24020201).

## Author contributions

A.B. and D.X. performed data analysis, plotted the manuscript figures, and drafted the manuscript. Y.L. and C.Y. collected wheat seeds and organized germplasm information from wheat populations around the world. A.B. performed the planting experiments with help of J.W.. A.B. performed read alignments with the help of J.Z.. L.K.,Y.G., X.S. and Z.Z. helped analyze the genomic data; X.Z., A.B. and D.X. performed the phylogenetic relationship. F.L. conceived the study and coordinated the project. All co-authors contributed to and edited the final version.

## Data availability

All 306 raw novel resequencing data were deposited at Genome Sequence Archive (GSA) under BioProject PRJCA001985 with BioSample subSAM090458 listing all accession numbers for individual samples.

## Code availability

We divide our code into two parts. All code to perform variants discovery are provided at Toolkit s Integrated for Genetic and Evolutionary Research (TIGER) https://github.com/PlantGeneticsLab/TIGER with -p FastCall. Other code to perform genomic analysis and regenerate all the figures in this manuscript is available at https://github.com/Fei-Lu/PrivatePlantGenetics/tree/master/src/analysis/wheat/VMap2.

## Supplementary Materials

Additional Materials and Methods

Supplementary Figures S1-S7

Supplementary Tables S1-S6

References (1-12)

