## Supplementary_materials_and_figures for "An integrated map of genetic variation from 1,062 wheat genomes"

**This Supplementary Information file includes:**

Materials and Methods

Supplementary Figures 1-7

Supplementary References

**Other Supplementary Materials for this manuscript include the following:**

Supplementary Table 1-6 as a separate Excel file

Materials and Methods

1 Overview of the collection of VMap 2.0 samples

1.1 Wheat germplasm collection and distribution

In this project, aiming to provide a comprehensive description of the large amount of common and rare variation in the wheat, we collected 1,125 accessions with different ploidy level and a broad set of continental backgrounds. The sample contains a total of 831 hexaploid wheat, 255 tetraploid wheats, as well as 39 *Ae. tauschii*, which are distributed in 85, 41 and 8 countries (Armenia, Azerbaijan, Iran, Russia, Soviet Union, Syria, Turkey, United States) relatively (Supplementary Fig. 5a). Among them, 306 samples are novel re-sequencing while the others are previously published ^1–5^. Detailed metadata information is available in Supplementary Table 1.

1.2 Germplasm repeatability check

To avoid duplicate of germplasm accessions, we have carried out a one-by-one inspection and found that there are 40 pairs of tetraploids as well 10 pairs of hexaploids with the same accession number. Combing the following initial genotype data, phylogenetic relationship and identity-by-state (IBS) distance will be performed to validate the duplicate samples. Finally, the qualified samples will be merged to produce unique one.

2 Data generation and processing

2.1 Whole-genome sequencing

The wheat population sampling plan was based on the goal of aggregating as much diversity as possible and calling rare mutations, so we complemented this with 306 newly re-sequencing samples consisting of 200 bread wheat, 80 emmer wheat and 26 *Ae. tauschii*. Genomic DNA was extracted from yellowed leaves at tillering stage and only high-quality DNA with content more than 1 μg and concentration more than 12.5 ng/μL can be used for library construction based on DNA NanoBalls (DNBs) technology. Then the whole-genome sequencing dataset was generated in two sequencing runs of paired-end 100 bp (PE100) by BGISEQ-500 sequencer based on combined primer anchor synthesis (cPAS). For hexaploid wheat, each sample library was sequenced on about 2 lanes, but for tetraploid and *Ae. tauschii*, each sample library was sequenced on only 1 lanes.

In summary, 469,864,946,476 raw reads were got, and genomes were sequenced at an average coverage of 10x with insert size 300 bp for newly resequencing individuals. The raw FASTQ files of the remaining 819 (1,125-306) accessions were obtained from publicly released data with coverage widely range from 3.8X to 34X, included part of the samples in pilot phase 1 (VMap 1.0 and VMap 1.1). Details on accessions count among different ploidy, average coverage, sequencing platform and paired-end size and insert size library were listed in Supplementary Table 2. We finally got a collection of 1,125 accessions with mean depth 9X.

2.2 Alignment and BAM processing

We perform the quality control of raw sequencing reads using SOAPnuke ^6^ before aligned. The adapter sequences were first filtered. Reads that contained “N” or quality value Q ≤ 15 accounting for more than 10% of the entire read were also filtered. FastQC were also used to check the random sampled clean reads. To create a comprehensive reference panel, clean reads were aligned to the wheat reference genome (IWGSC RefSeq v1.0) using Burrows Wheeler Aligner - MEM (Version: 0.7.17-r1188) for each sequence lane. It is particularly worth noting that samples with different polyploid would be aligned to the consistent subgenome of reference Chinese Spring (CS) as described previously (e.g. tetraploid would be mapped to A and B subgenome of reference genome) ^7^. Initial SAM file went through the following process using SAMtools ^8^: name-sorted, mate information fixed, coordinate-sorted and duplicate reads removal.

SOAPnuke1.5.6 filter -f AAGTCGGAGGCCAAGCGGTCTTAGGAAGACAA -r AAGTCGGATCGTAGCCATGTCGTTCT

GTGAGCCAAGGAGTTG -1 $raw_1.fq.gz -2 $raw_2.fq.gz -C $1.clean.fq.gz -D $2.clean.fq.gz -o $outputPath -l 15 -q 0.1 -n 0.1 -Q 2 -G -5 0

bwa mem -t 8 -R '@RG\tID:inputName\tPL:illumina\tSM:inputName\tLB:inputName' wheat_iwgscV1.fa.gz input_1.clean.fq.gz input_2.clean.fq.gz | samtools view -S -b - > $output.pe.bam && echo "** bwa mapping done **" &

samtools sort -n -m 20G -o $output.namesort.bam -O bam -@ 40 $input.pe.bam

samtools fixmate -m $input.namesort.bam $output.fixmate.bam

samtools sort -m 20G -o $output.fixmate.pos.bam -O bam -@ 40 $input.fixmate.bam

samtools markdup -r $input.fixmate.pos.bam $output.rmdup.bam

samtools index $input.rmdup.bam

2.3 BAM quality control

To review the success of alignment and de-duplication, we performed several BAM qualities control. We random sampled the final BAM files and then checked if adapter sequences, possible contaminations and duplicates have been marked or removed successfully. To enhance the power of variation discovery, reads coverage depth, insert size and GC content will also be assessed in this processing.

samtools stats -r wheat_iwgscV1.fa. $input.rmdup.bam > $output.rmdup.bam.bc

perl $samtools-1.8/misc/plot-bamstats -p $input.rmdup.bam.bc

2.4 Overview of data processing pipeline for VMap 2.0

We initially collected 1,125 diverse individual genomes for whole-genome resequencing with the goal of evaluating the dynamic evolution of mutation burden among hexaploid wheat populations and its wheat relatives. Here we listed a summary pipeline of steps that went into building the second-generation genetic variation map (VMap2.0). First, to ensure a consistent coordinate system for variant sites, each sequencing dataset was uniformly aligned to the reference genome IWGSC RefSeq v1.0 and was processed by a standardized custom pipeline (to perform the SNP discovery and genotyping). Second, considering the enriched false positives variants probably caused by false read mapping and genotyping ^9^, ‘accessible reliable reference genome’ was needed to remove these artefacts. Based on our newly developed PopDep (<https://github.com/PlantGeneticsLab/TIGER/wiki/PopDep>), we identified the conserved/single-copy/well-assembled region and filtered regions with extent plenty of ambiguously placed reads and unbelievably high or low alignments from the reference genome, and then kept the intersection of reliable sites from different ploidy wheat, we finally obtained a high-quality reliable genome database (RGDB) with reliable alignment to the correct genome location. The RGDB was used to filter the initial single nucleotide variant (SNV) and short (≤50bp) insertion/deletion (indel) for a relatively high standard. Third, we genotyped all individuals using FastCall2 pipeline and further carried out other data filtering operations: variants with site heterozygosity ≤ 0.05, site missing rate ≤ 0.2 and minor allele occurrence (MAC) ≥ 2 were kept generating a comprehensive variation map. The final dataset contains 1,062 taxa, 196 millions bi-allelic single nucleotide polymorphisms (SNPs), 2.2 millions insertions and 4.7 millions deletions, with the average depth of 9X.

2.5 Sample provenance check and removal for finalizing samples 1,062 final accessions

To check the repeatability of the sample, the pairwise identity by state (IBS) distance were computed using TASSEL 5 ^10^. In total, 49 duplicates and 1 triplicate were identified as the same taxa and showed a consistent phylogenetic tree structure. However 13 taxa (1.84%) were removed due to the abnormal cluster on phylogenetic tree and principal component analysis, leaving a dataset of 1,062 samples for VMap2.0 finalization. The removed samples are as follows: IG140057, PI583718, PI534284, Beaqle, and Caruton for hexaploid, PI560877, PI300990, PI466930, PI466959, PI272522 for tetraploid, as well as KU-2071, TA2462, AE430 for *Aegilops tauschii*. The remaining 1,062 taxa were clustered into different genetic groups: *Aegilops tauschii*, Wild emmer, Domesticated emmer, Free-threshing tetraploids, Landrace, Cultivar or Other.

3 Variants effect prediction algorithms

VEP and SnpEff are critical tools for annotating and prioritizing genetic sequences in whole genome sequence study, which can systematically analysis most types of genomic variation and return detailed annotation on the effect of different transcript isoforms as well as regulatory regions. Typically, four categories of impact were included in the final output: modifier, low, moderate and high. However, both approaches rely on the genome assembly quality. Variants with high effect and relatively lower number are most located in biological unit of start_lost, stop_gain/lost, splice_accepor/donor region, which will be considered as putatively deleterious.

4 Analysis

4.1 Phylogenetic tree construction

To figure out the evolutionary history of bread wheat and its wild ancestors, we performed the stepwise phylogenetic analysis. First, barley was used as an outgroup to identify the most recent common ancestors of wild emmer. A subset of 34,383 SNPs that could be uniquely aligned to the barley genome (assembly version: IBSC_v2) was selected to construct the phylogenetic tree by RAxML (v 8.2.12). The command line was raxml -f a -m PROTGAMMAWAG -p 12345 -x 12345 -# 100 -s Asubgenome_addbarley.phy -n Asubgenome_addbarley.raxml -o barley -T 80. And then, the second tree was reconstructed based upon ~230,000 bi-SNPs on AB subgenomes of VMap 2.0 with MAF > 0.1, which included 1,026 accessions. And the command line was raxml -f a -m GTRGAMMA --JC69 -p 12345 -x 12345 -# 100 -s ABsubgenome_01.phy -n ABsubgenome_01.raxml -o out -T 100. Finally, the newick format results were visualized using iTOL^11^ (<https://itol.embl.de/>).

4.2 Population genetics parameters

The genome-wide nucleotide diversity (π) of different wheat populations and population differentiation index (*F*_ST_) were calculated using VCFtools v0.1.15 ^12^ software. The command was as followings:

vcftools --gzvcf chr1A_vmap2.1.vcf.gz \

--keep 000_group/WE.txt \

--window-pi 100000 --window-pi-step 50000 \

--out WE_chr1A_based100000Window_50000step

vcftools --gzvcf chr1A_vmap2.1.vcf.gz \

--weir-fst-pop ../000_group/hexa_tetra/LR_AM.txt \

--weir-fst-pop ../000_group/hexa_tetra/LR_CSA.txt \

--fst-window-size 100000 --fst-window-step 50000 \

--out LR_AM_VS_LR_CSA_chr1A

4.3 Population structure analysis

To perform principal component analysis (PCA), we randomly selected 500 kb SNPs from the entire genome. We then used TASSEL 5 software ^10^ to generate an Identity-by-state (IBS) matrix. Finally, we conducted dimensionality reduction analysis using the PCA function in the R packages factoextra and FactoMineR, and visualized the first two principal components.

4.4 Statistical test

We performed a preliminary Shapiro-Wilk test to make sure that the two groups being compared are normally distributed. And then performed the Levene’s test to check the homogeneity of variances. Other details were described in the methods.

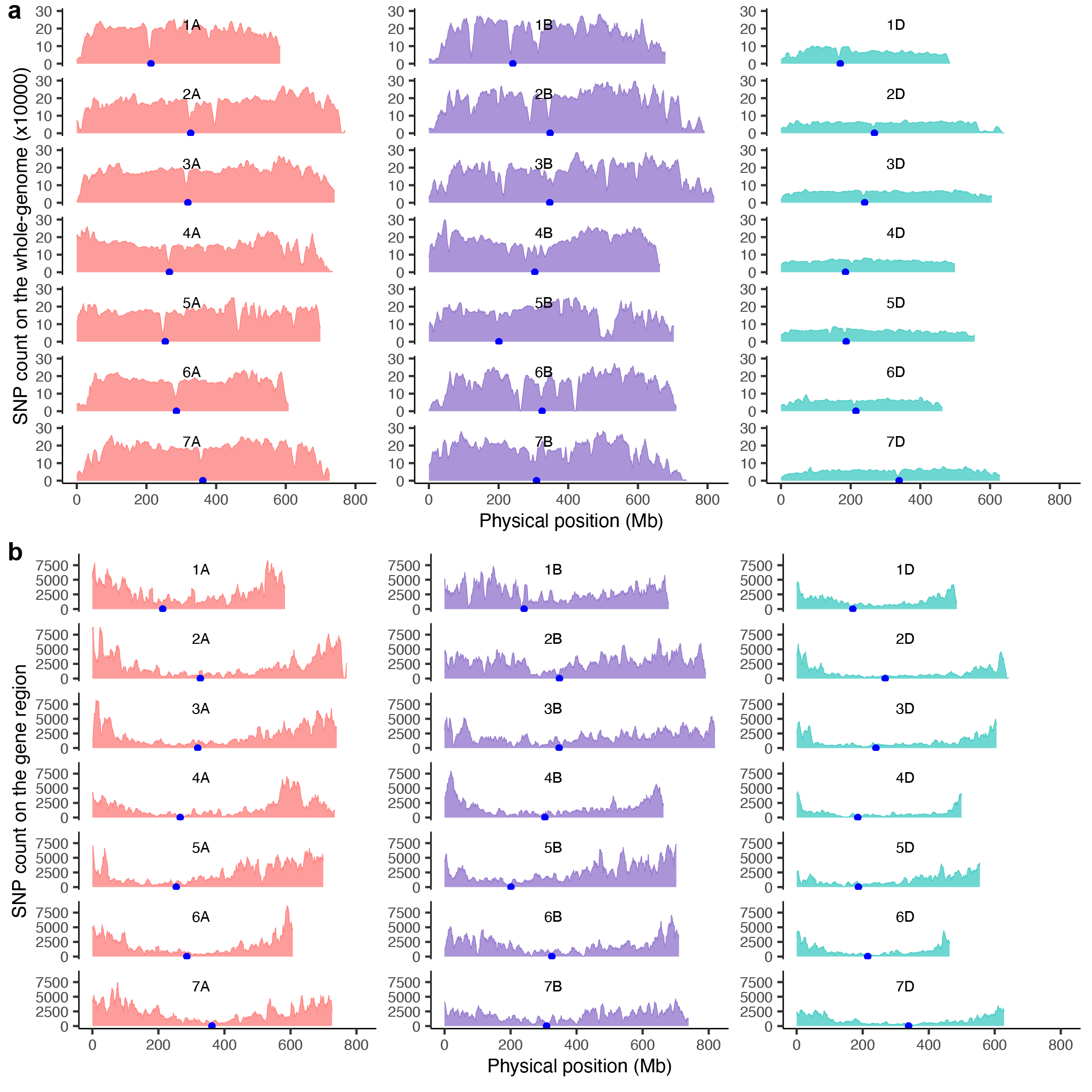

Supplementary Fig. 1 | The distribution of polymorphic SNPs in VMap 2.0 dataset.

**a,** whole genome regions. **b,** gene regions.

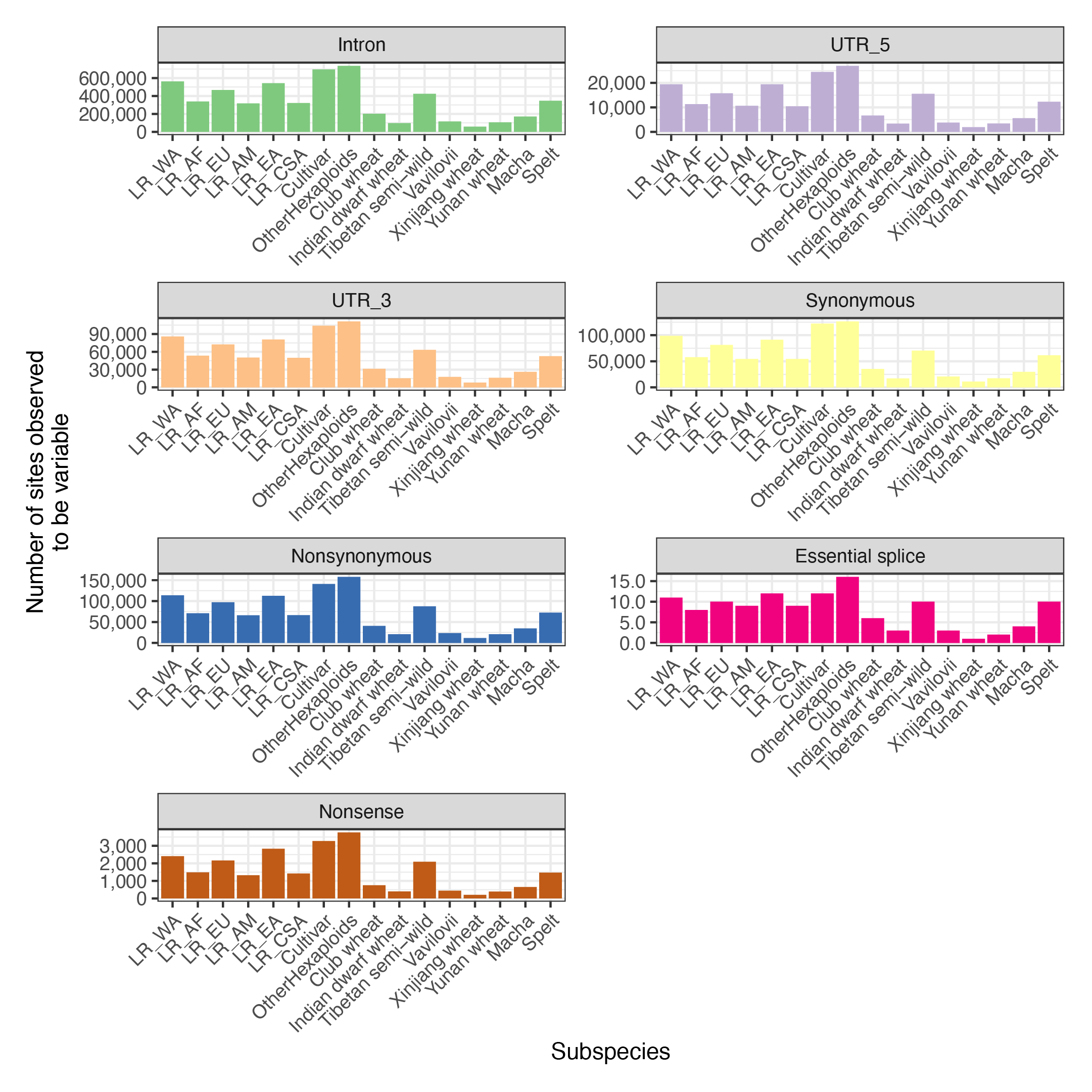

Supplementary Fig. 2 | Number of SNPs in different subpopulations of hexaploid wheat.

The types of variants in this study mainly include introns, 5'UTRs, 3'UTRs, synonymous mutations, nonsynonymous mutations, essential splice sites, and nonsense mutations in gene regions.

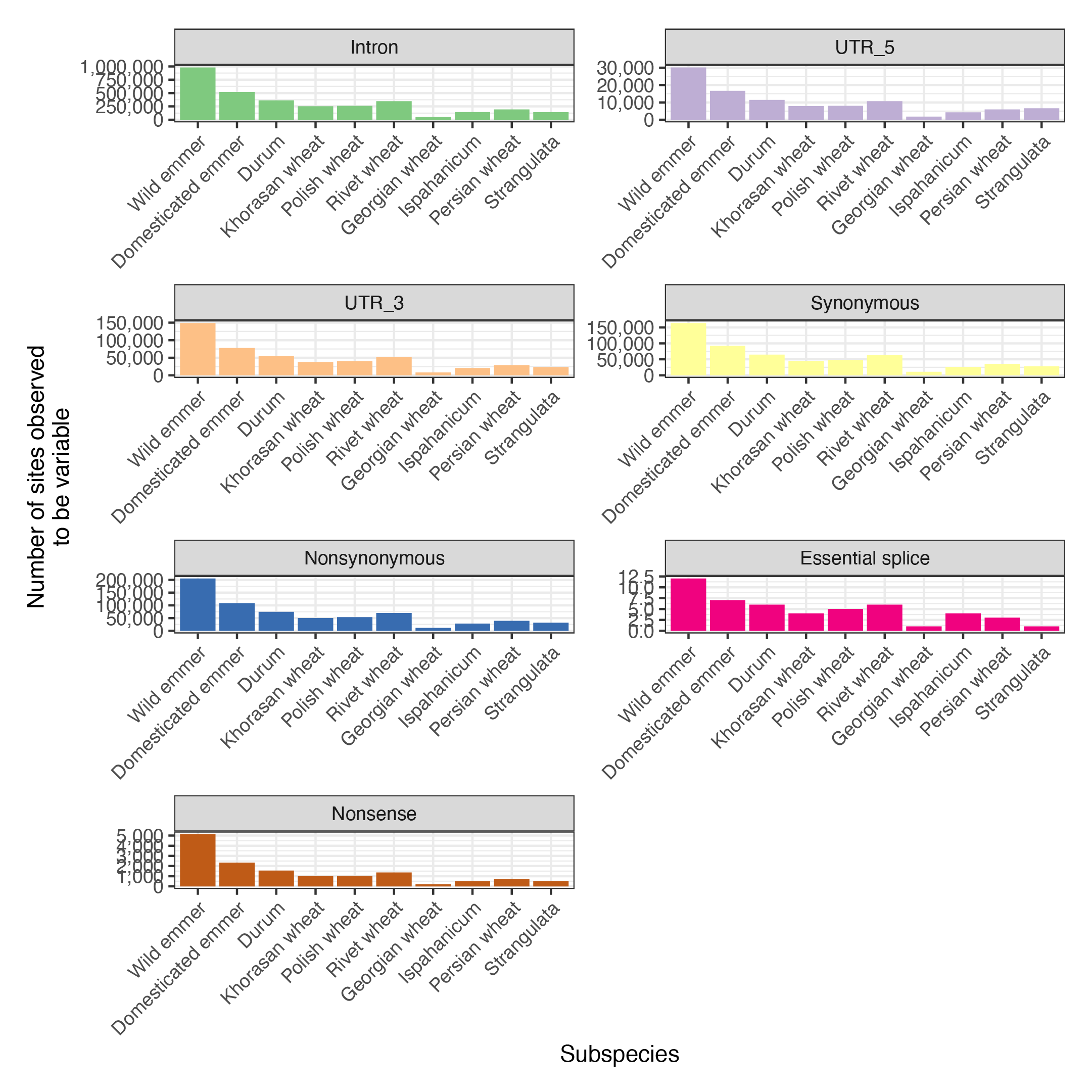

Supplementary Fig. 3 | Number of SNPs in diploid *Ae.tauschii* and tetraploid wheat subpopulations.

The types of variants in this study mainly include introns, 5'UTRs, 3'UTRs, synonymous mutations, nonsynonymous mutations, essential splice sites, and nonsense mutations in gene regions.

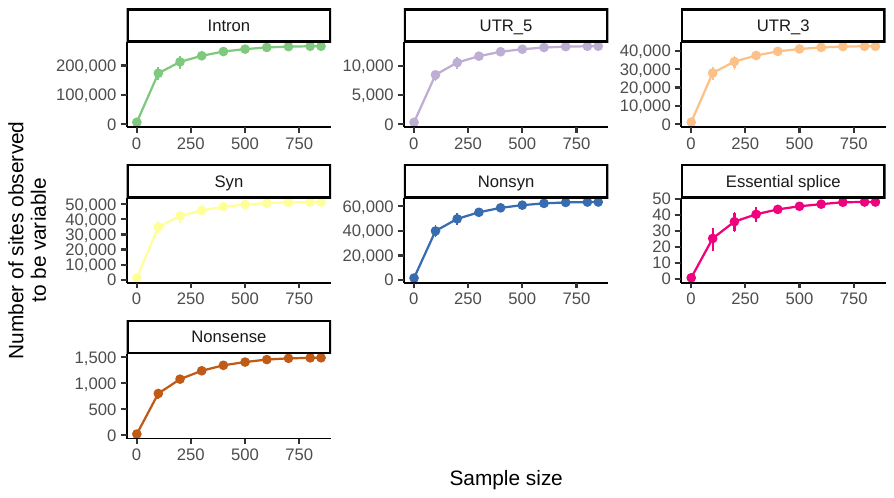

Supplementary Fig. 4 | The total number of variants observed as a function of sample size in D subgenome.

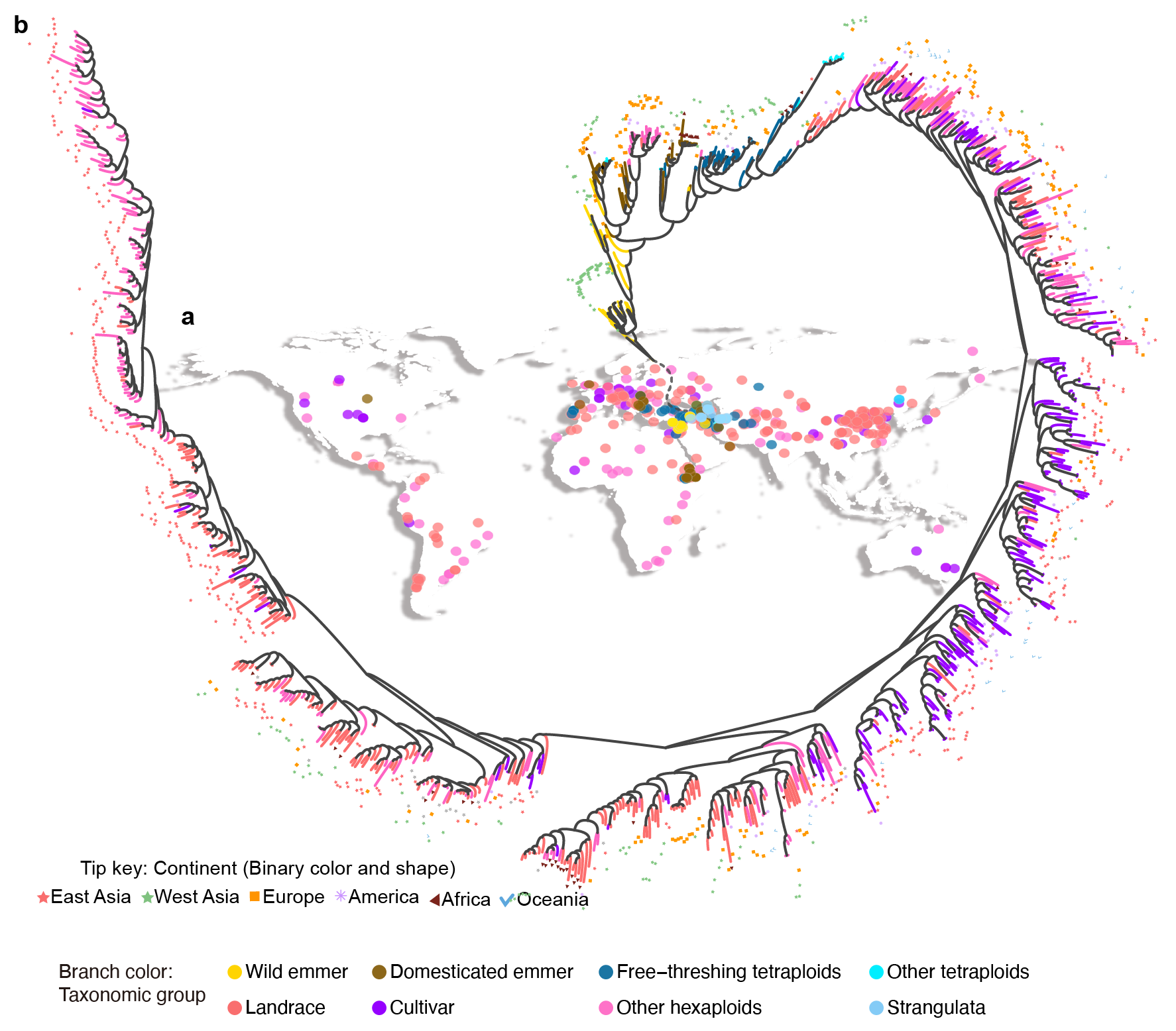

Supplementary Fig. 5 | Population analysis of 1062 wheats.

**a,** Geographical distribution of wheat and its wild relatives samples. **b,** The phylogenetic relationships across tetraploids wheat and hexaploid wheat.

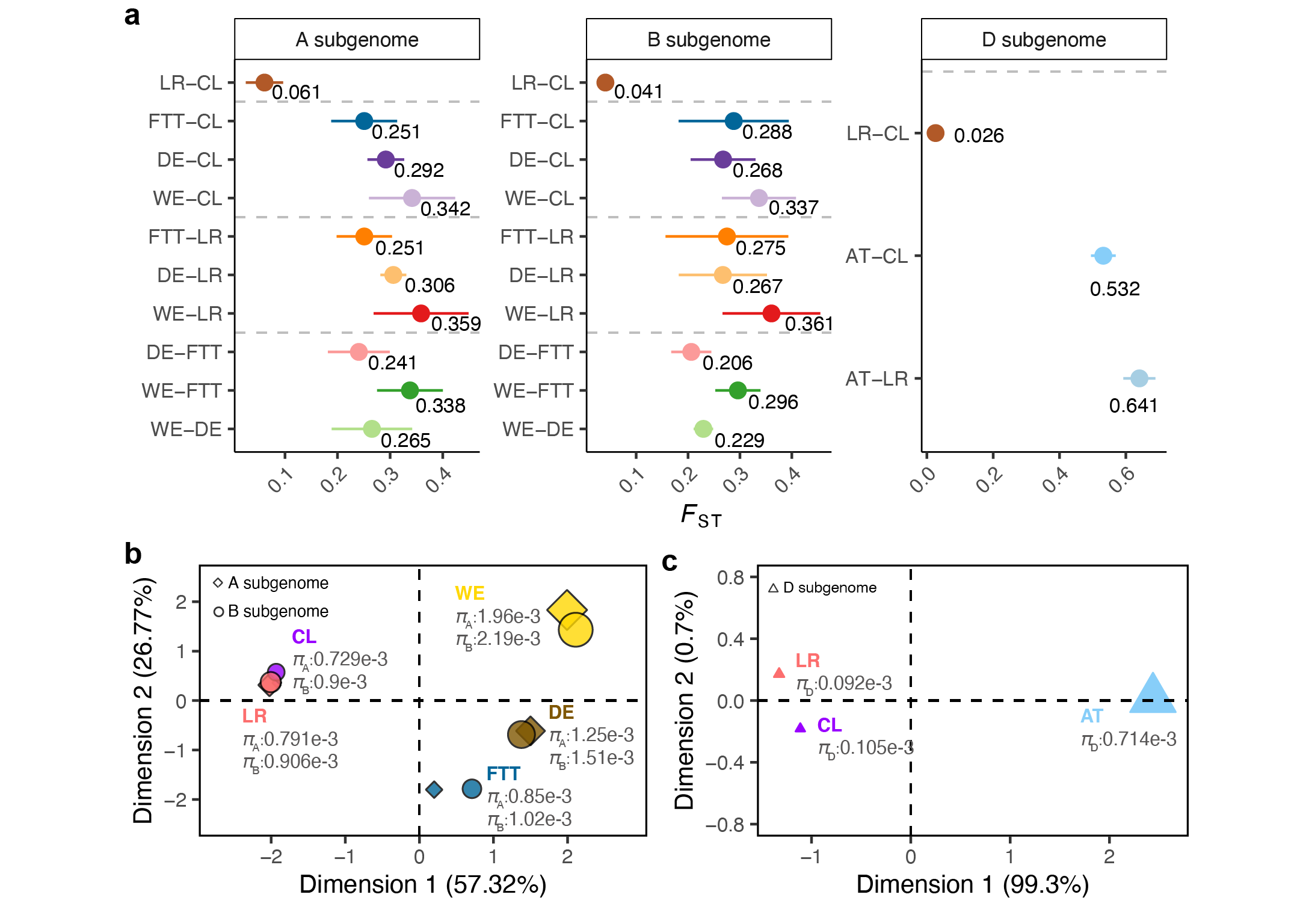

Supplementary Fig. 6 | Genetic differentiation of the different wheat species.

a, Pointrange plot showing population differentiation (*F*_ST_) among diverging populations. Points denote the mean value and error bars represent 95% confidence intervals (sometimes error bars are fully contained within the point). b-c, Multidimensional scaling of pairwise *F*_ST_ values between different subpopulations in AB subgenomes (b) and D subgenome (c). The size of different shapes represents the nucleotide diversity (*π*). WE, Wild emmer; DE, Domesticated emmer; FTT, Fresh-threshing tetraploids; LR, Landrace; CL, Cultivar; AT, *Ae. tauschii*.

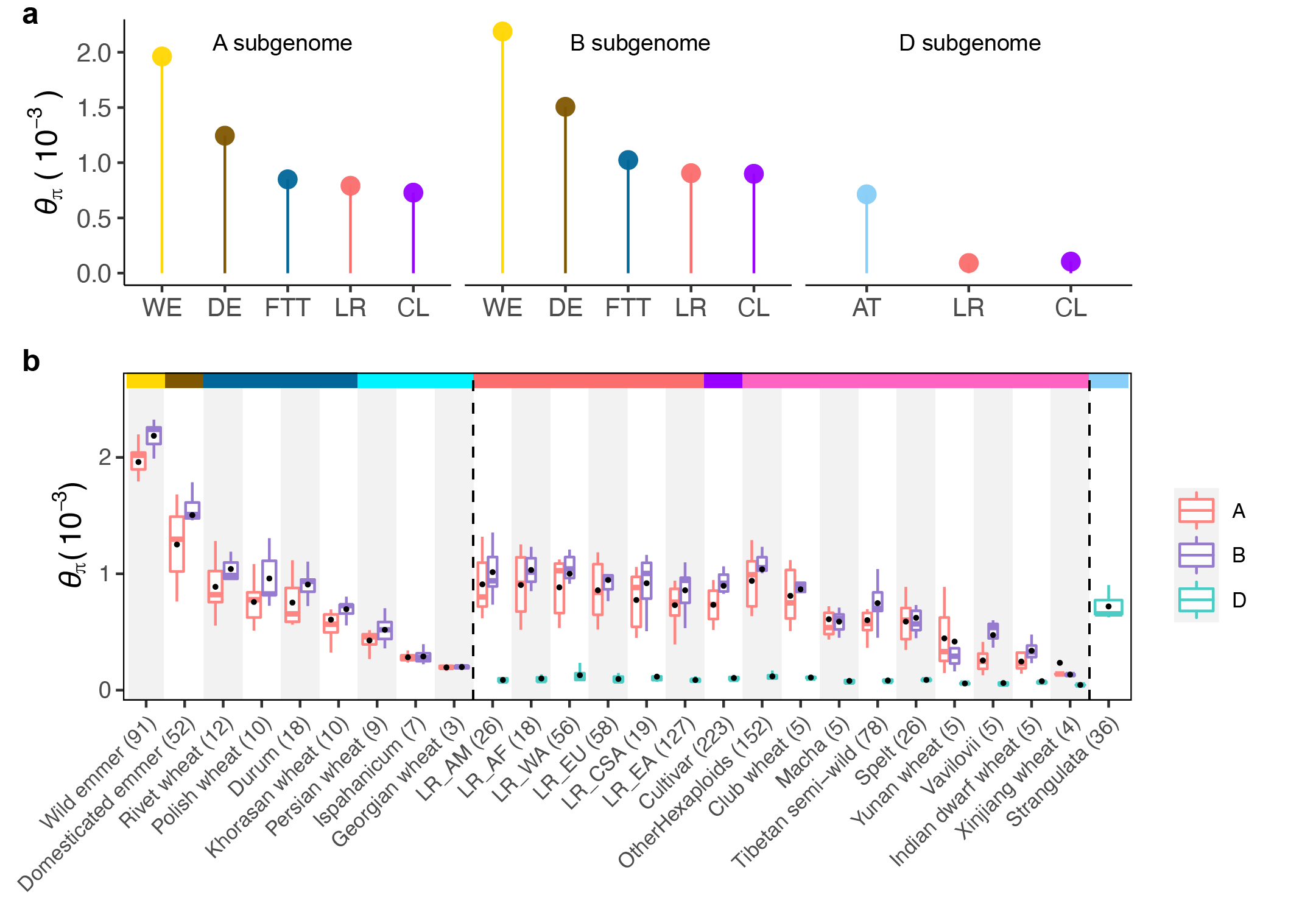

Supplementary Fig. 7 | Genome-wide mean nucleotide diversity in different wheat subpopulations.

**a,** Average nucleotide diversity (*θ*_π_). WE: Wild emmer; DE: Domesticated emmer; FTT: Free-threshing tetraploids; LR: Landrace; CL: Cultivar; AT: *Ae. tauschii*. **b,** Nucleotide diversity in all subpopulations grouped by subgenomes (*n* = 1,062). The centerline of each box plot indicates the median and the lower and upper hinges indicate the 25th and 75th percentiles, respectively. The vertical line of each boxplot extends to 1.5× the interquartile range from each hinge. The black dot in the middle of the boxplot represents the mean value. The striped rectangle above the figure represents the largest subpopulations classification, and the colors are consistent in **(a)**.
